# Phage-Derived Depolymerase as an Antibiotic Adjuvant Against Multidrug-Resistant *Acinetobacter Baumannii*

**DOI:** 10.1101/2021.05.26.445908

**Authors:** Xi Chen, Miao Liu, Pengfei Zhang, Miao Xu, Weihao Yuan, Liming Bian, Yannan Liu, Jiang Xia, Sharon S.Y. Leung

## Abstract

Bacteriophage-encoded depolymerases are responsible for degrading capsular polysaccharides (CPS), lipopolysaccharides (LPS) and exopolysachcharides (EPS) of the host bacteria during phage invasion. They have been considered as promising antivirulence agents in controlling bacterial infections, including those caused by drug-resistant bacteria. This feature inspires a hope of utilizing these enzymes to disarm the polysaccharide capsid of the bacterial cells, which then strengthens the action of antibiotics. Here we have identified, cloned, and expressed a depolymerase Dpo71 from a bacteriophage specific for the gram-negative (G-ve) bacterium *Acinetobacter baumannii* in the heterologous host *E. coli*. Dpo71 sensitizes the multidrug-resistant (MDR) *A. baumannii* to the host immune attack, and also acts as an adjuvant to assist or boost the action of antibiotics, for example colistin. Specifically, Dpo71 at 10 µg/ml enables a complete bacterial eradication by human serum at 50% volume ratio. Dpo71 inhibits biofilm formation and disrupts the pre-formed biofilm. Combination of Dpo71 could significantly enhance the antibiofilm activity of colistin, and improve the survival rate of *A. baumannii* infected *Galleria mellonella*. Dpo71 retains the strain-specificity of the parent phage from which Dpo71 is derived: the phage-sensitive *A. baumannii* strains respond to Dpo71 treatment, whereas the phage-insensitive strains do not. This indicates that Dpo71 indeed is responsible for the host specificity of bacteriophages. In summary, our work demonstrates the feasibility of using recombinant depolymerases as an antibiotic adjuvants to supplement the development of new antibacterials and to battle against MDR pathogens.

## INTRODUCTION

Carbapenem-resistant *Acinetobacter baumannii* was identified as number one priority pathogen by the World Health Organization (WHO) in 2017 [1, 2]. *A. baumannii* infection is associated with frequent and hard-to-treat infections, such as pneumonia, bacteremia, urinary tract infections, meningitis and wound infections [3]. In the past-decades, outbreaks of *A. baumannii* resistant to the last-resort antibiotics, such as colistin, are increasingly reported [4, 5]. Alarmingly, the development of novel antibiotics have experienced significnat setbacks in recent years [6], and thereby the field is urgenly calling for novel antibacterial agents to address the clinical challenges of *A. baumannii* associated infections.

Bacteriophages (phages), natural co-evolving bacteria killers, are being revitalized to combat multidrug-resistant (MDR) bacteria [7]. Although regarded as a promising alternative to conventional antibiotics, phage therapy faces challenges as all the completed clinical trials failed to confirm the efficacy. The narrow host range and the development of phage-resistance could be the major factors for such failures. The viral nature of phage may also be unacceptable to most clinicians and the general public [8‒10]. Alternatively, researchers explore the potential of phage-encoded enzymes, including the peptidoglycan hydrolases (PGH), polysaccharide depolymerases, and holins, as novel antibacterial agents, drawing inspiration from the life cycle of phage. In contrast to phages, phage-encoded enzymes, as therapeutic proteins without replicating capability are more manageable and acceptable [11‒14].

Phage-encoded depolymerases are polysaccharide hydrolases or lyases responsible for stripping bacterial polysaccharides, including exopolysaccharides (EPS), capsular polysaccharides (CPS), and lipopolysaccharide (LPS), to facilitate the parent phage to inject its DNA materials into the bacterial host [13, 14]. Distinct from other phage-encoded enzymes, depolymerases do not lyse bacterial cells directly. Instead, they disintegrate the CPS of bacteria to make them susceptible to host immune attack and antibacterial treatment [15]. Recombinant depolymerases have been shown to protect mice from fatal systemic bacterial infection [16, 17] and to disrupt biofilms for enhanced antimicrobial activity [18‒22].

Combined administration of depolymerases and antibiotics will produce superior antibacterial efficacy – expected but not yet well-supported by experiments. Bansal *et al*. were the first to report the synergistic use of a depolymerase derived from *Aeromonas punctata* (a facultative anaerobic G-ve bacterium) with gentamicin in treating mice infected with non-lethal dose of *K. pneumonia* [23]. Intranasal administration and intravenous administration of the combination for lung infection and systemic infection, respectively, both reduced bacterial counts significantly more than the single-agent treatments. They attributed the improved bacterial killing efficiency to the enhanced bacterial susceptibility towards gentamicin after the bacteria were decapsulated by the depolymerase. Depolymerases also effectively dispersed the EPS matrix in *K. pneumonia* biofilms to facilitate the penetration of gentamicin [24]. A similar synergy was also observed in the treatment of *K. pneumonia* biofilms using the Dep42 depolymerase and polymixn B [25]. On the contrary, Latka *et al*. showed that the KP34p57 depolymerase had no impact on the activity of ciprofloxacin, but could significantly enhanced the antibiofilm efficiency of non-depolymerase-producing phages [26]. Depolymerase could also enhance the antibiofilm efficacy of a phage-encoded antibacterial enzyme endolysin [27].

Currently, a few *A. baumannii* depolymerases have been identified [20, 28‒32]. Whether they would work synergistically with antibiotics in controlling biofilm-associated infections, like those observed for *K. pneumonia*, remains questionable. In the present study, the combinational effects of a depolymerase Dpo71 encoded by a lytic *A. baumannii* phage, vB_AbaM-IME-AB2 (IME-AB2 in short), [33] with serum or colistin in targeting MDR *A. baumannii* are evaluated for the first time.

## RESULTS

### Identification and characterization depolymerase Dpo71

The IME-AB2 phage exhibits plaque surrounded by a halo-zone, suggesting the presence of a depolymerase protein. Bioinformatic analysis reveals that gp71 is a tailspike protein [34] with 43% sequence similarity as the depolymerase encoded by another *Acinetobacter* phage vB_AbaP_AS12 (Protein Data Bank number 6EU4) (**Fig. 1A**). Expression of the ORF71 sequence in *E. coli*. yields a protein with more than 95% purity and a molecular mass of about 80 kDa (**Fig. 1B**), matching the calculated value of 80.2 kDa. Size exclusion chromatography shows that the purified Dpo71 elutes as a single peak at a molecular weight larger than 200 kDa (**Fig. 1C**). This corresponds to a trimer, consistent with the expected oligomeric form of phage tail fiber protein which is believed to endure extreme conditions for phage infection and survival [30, 35]. Circular dichroism (CD) reveals that the Dpo71 protein adopts a well-folded conformation rich in β-sheet structures with a negative dichroic minimum at 215-nm and a positive maximum around 195-nm characteristic peaks (**Fig. 1D**). The melting curves following the CD signal at 215 nm show a melting temperature (Tm) of 58.5 °C (**Fig. 1E**). Spot tests were performed next to confirm the ability of Dpo71 in degrading bacterial capsules of the host bacteria MDR-AB2 of the parent phage. Semi-clear spot formation was observed on the bacterial lawn with the spot sizes increasing with the dose of depolymerase from 0.001 µg to 10 µg range (**Fig. 1F**). The EPS degradation activity of Dpo71 was also confirmed by the reduced EPS concentration and Alcian blue staining (**Fig. S1**) using protocols documented in previous studies [36, 37].

**FIG 1.**
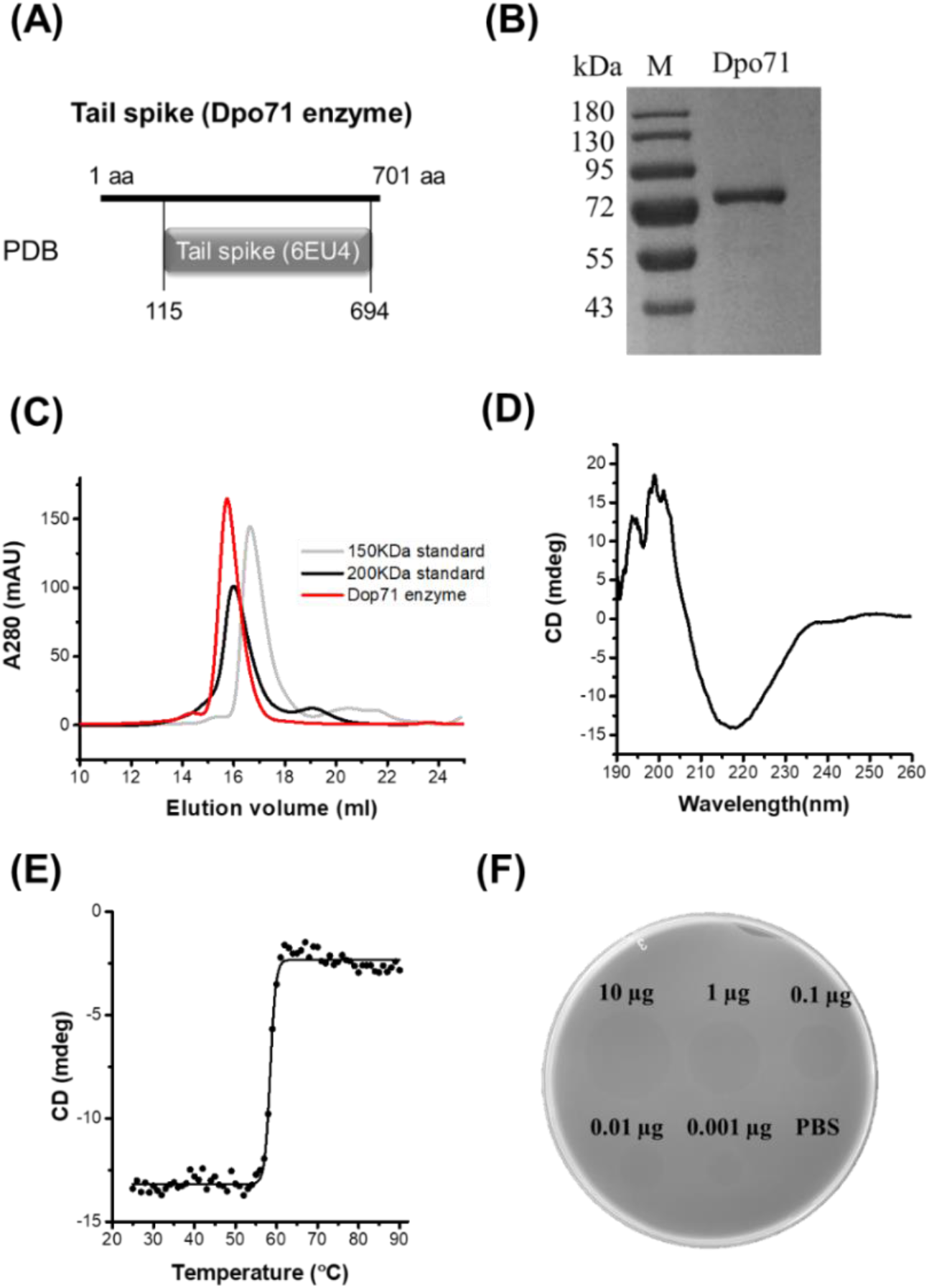
Identification and characterization of Dpo71 depolymerase. (A) Bioinformatic analysis indicates the gp71 gene of the IME-AB2 phage. (B) SDS gel electrophoresis analysis of purified Dpo71 and a standard molecular mass marker (Lane M). (C) Size exclusion chromatography of purified depolymerase protein. (D) Circular dichroism analysis of Dpo71 measured in the far-UV (190 to 260 nm). (E) Melting curve of Dpo71 acquired at 215 nm from 20 to 90°C. (F) Spot test assay of Dpo71 against *A. baumannii* lawn (0.001µg to 10 µg).

### Robustness of Dpo71 upon administration and storage

To evaluate the therapeutic applicability of Dpo71, the enzymatic activity of Dpo71 at various pH was evaluated by monitoring the turbidity of the residual EPS. **Fig. 2A** shows the enzyme remained active in the range of pH 4 to 8, covering most of the physiological conditions. Then we measured the toxicity of Dpo71 to mammalian cells, human red blood cells and lung bronchial epithelial cell line (BESA-2B cells). No haemolytic activity was detected even at a high dose of 500 µg /ml (**Fig. 2B**). **Fig. 2C** also shows Dpo71 has no cytotoxicity against BESA-2B cells. The result suggests that Dpo71 may be a safe treatment, likely for systemic or pulmonary infections. As stability upon storage is critical for the development of commercially viable protein therapeutics, the storage stability of Dpo71 at 4 °C has been evaluated. Results show that Dpo71 is stable for at least 6 months without noticeable activity loss (**Fig. 2D**).

**FIG 2.**
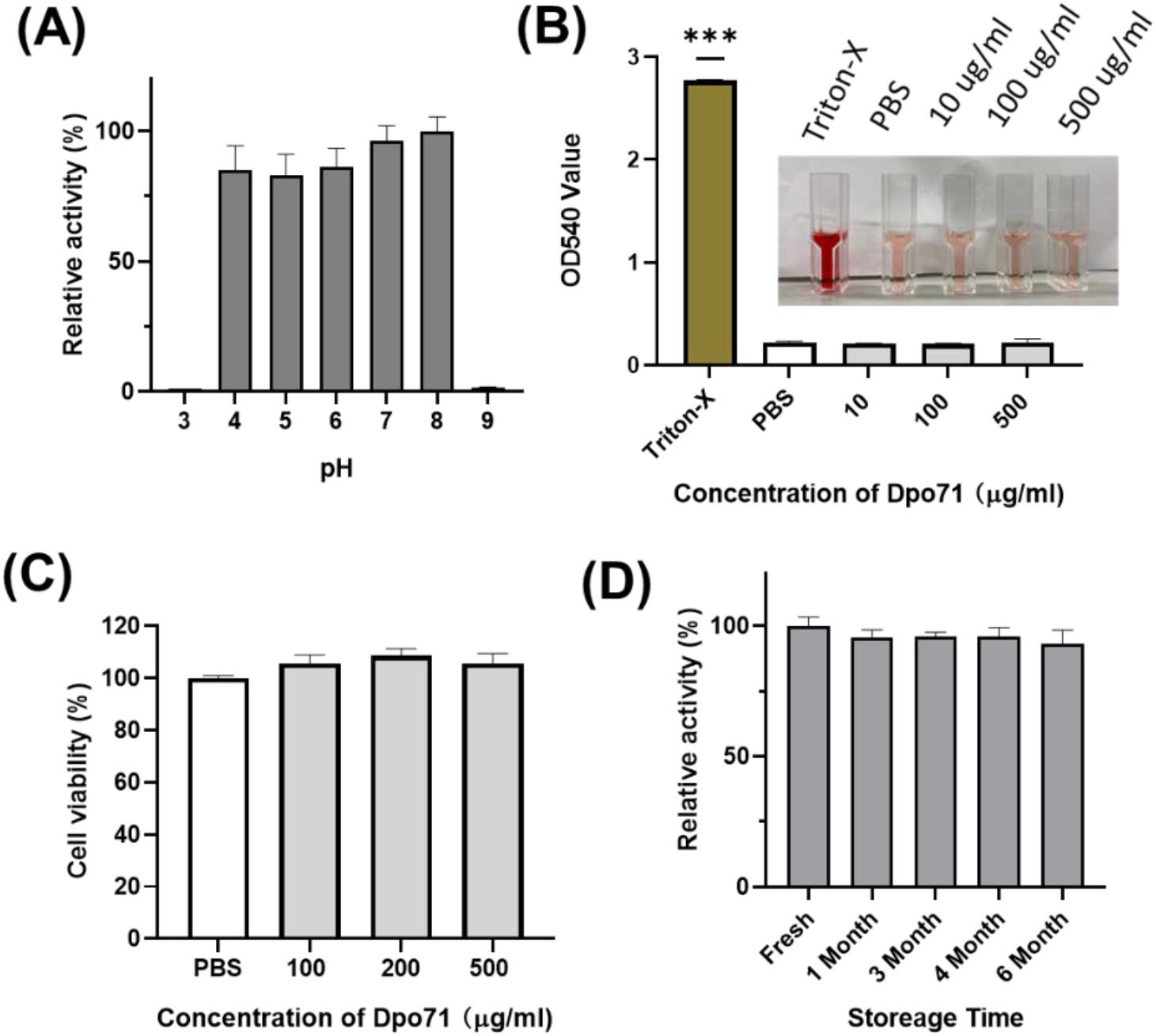
Stability and toxicity of Dpo71. (A) The effect of pH on Dpo71 depolymerase activity. CPC turbidity assay was used to measure the EPS degradation activity, the highest activity at pH 8 was set as 100%. (B) Hemolysis of red blood cell by Dpo71. PBS and 0.1% Triton X-100 in PBS as the negative and positive controls, respectively. *** indicated P < 0.001. (C) Cytotoxicity of Dpo71 against human cells (BESA-2B). (D) Stability of Dpo71 after storing at 4 °C for specific month. The EPS degradation activity of fresh opened enzyme was set as 100%. Data are expressed as means ± SD (n = 5).

One of the limitations of phage therapy is known to be the development of resistance. So, we next evaluated whether Dpo71 can induce resistance development. The MDR-AB2 strain was challenged with the parent IME-AB2 phage, Dpo71 or PBS for 24 h and tested for the emergence of resistant phenotypes (**Table 1**). Ten bacterial colonies of each culture were selected to estimate the sensitivity to the phage or Dpo71 using the spot test assay. Representative spot test figures are shown in **Fig. S2**. PBS treated bacteria all remained sensitive to both the IME-AB2 phage and Dpo71. Pre-challenged with IME-AB2 phage resulted in resistance to both the phage and Dpo71 afterwards. Although Dpo71 was originated from the same phage, all ten bacterial colonies tested remained susceptible to the enzyme without resistance development. This observation also agrees with the findings reported by *Oliveira* et al. [31, 37]. These data, although preliminary, show that Dpo71 as a therapeutic option is less likely to induce resistance than phages.

**Table 1.** Bacterial resistance table of depolymerase Dpo71 on *A. baumannii* (+ indicating sensitive; - indicating insensitive).

| No. of isolates | Incubation with PBS |  | Incubation with Phage |  | Incubation with Dpo71 |  |
| --- | --- | --- | --- | --- | --- | --- |
|  | Phage | Dpo71 | Phage | Dpo71 | Phage | Dpo71 |
| 1 | + | + | – | – | + | + |
| 2 | + | + | – | – | + | + |
| 3 | + | + | – | – | + | + |
| 4 | + | + | – | – | + | + |
| 5 | + | + | – | – | + | + |
| 6 | + | + | – | – | + | + |
| 7 | + | + | – | – | + | + |
| 8 | + | + | – | – | + | + |
| 9 | + | + | – | – | + | + |
| 10 | + | + | – | – | + | + |

### Sensitizing bacteria to serum killing and antibiotic

As depolymerase can disintegrate bacterial capsules and thus sensitize bacteria to kill by host immune system [35, 38, 39], we first measured whether serum could kill the Dpo71-treated bacterial cells. Two *A. baumannii* strains (AB#1 and AB#2) sensitive to the parent IME-AB2 phage and other two insensitive (AB#3 and AB#4) strains were chosen for the serum killing assays (**Table S1**). The four tested strains were resistant to serum killing and continue to grow in human serum without the presence of depolymerase. Bacteria treated with 10 µg/ml Dpo71 and inactivated serum also showed no antibacterial effect (**Fig. 3A**). On the contrary, a remarkable bacterial reduction (8 log) was noted for the two sensitive strains (AB#1 and AB#2), but not for the insensitive ones (AB#3 and AB#4) when the bacteria were treated with active human serum (with a volume ratio of 50%) and Dpo71 (**Fig. 3A**). Furthermore, the time killing assay on the two sensitive strains was performed with a serum ratio of 1% to 50%. Complete bacterial eradication was observed after a five-hour treatment at 50% human serum for both AB#1 and AB#2 (**Fig. 3B** and **S3**). It is noteworthy that a 5% serum was sufficient to achieve around 4-log bacterial reduction after 5 h and with minor regrowth after 24 h. This efficacy is significantly higher as compared with previous reports, in which at least 25% human serum is needed to achieve the same killing efficiency [15, 21, 31, 35, 36].

**FIG 3.**
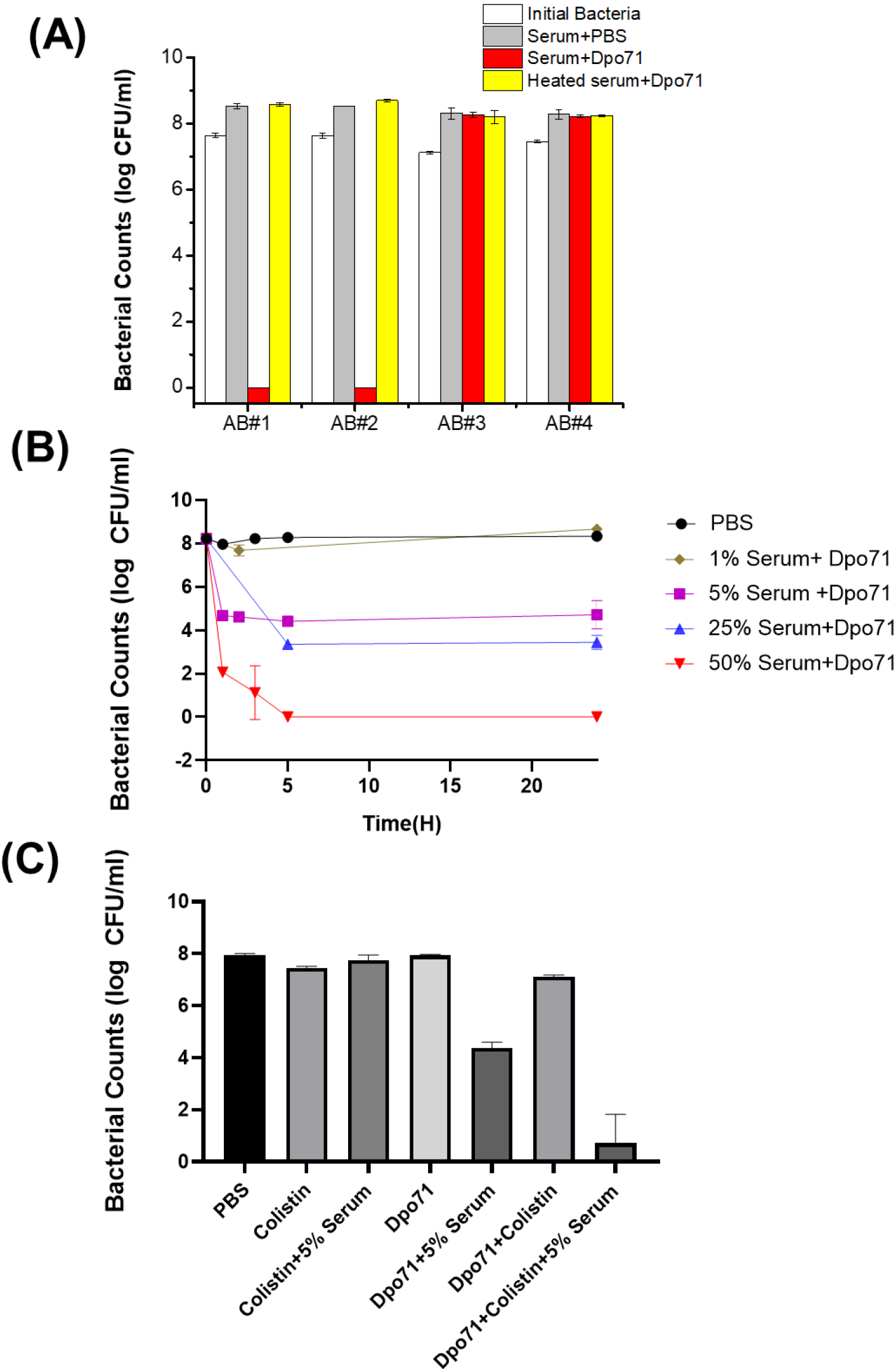
Dpo71 enhanced the serum sensitivity and colistin activity against *A. baumannii*. (A) Bacterial susceptibility to the treatment with Dpo71 concentration of 10 µg/ml and 50% serum volume ratio. *A. baumannii* clinical strains sensitive (AB#1 and AB#2) and insensitive (AB#3 and AB#4) to the phage IME-AB2 were tested. (B) Time-killing curve of Dpo71 (10 µg/ml) against MDR-AB2 (AB#2 strain) in the presence of 1% to 50% human serum. (C) Dpo71 and colistin activity against *A. baumannii* in the presence / absence of 5% human serum. Data are expressed as means ± SD (n = 3).

We next examined the antibiotic adjuvant effect of Dpo71. Colistin was chosen because it was the only tested antibiotics that the MDR-AB2 strain was susceptible to with a minimum inhibition concentration (MIC) of 2 μg/ml [34] (**Table S2**). The colistin concentration was set at 1 μg/ml, half of the MIC value. **Fig. 3C** shows that 1 ug/ml colistin alone or colistin with 5% serum had no antibacterial effect against the inoculation of 10^8^ cfu/ml. Dpo71 combined with 5% serum could achieve around 4-log bacterial reduction and the antibacterial effect was further enhanced to nearly complete eradiation (residual viable bacteria reduced from 4.4 ± 0.2 log to 0.7± 1.1 log) when Dpo71 was used in combination with 1ug/ml colistin in the presence of 5% serum (**Fig. 3C**). This boosting effect was consistent with a modified checkerboard assay examining the synergy between Dpo71 and colistin that the MIC of colistin dropped from 2 μg/ml to 0.5 μg/ml with the addition of 5% serum (Data not shown). Notably, these results indicated that Dpo71 could act as an adjuvant to boost the antibacterial activity of colistin in low serum condition.

### Anti-biofilm activity

We next measured the inhibition effect of Dpo71 on biofilm formation. **Fig. 4A** shows Dpo71 inhibits biofilm formation in a dose-dependent manner. At 1 μg/ml Dpo71, the residual biomass was 80% as compared with the PBS-treated control. The residual biomass was further reduced to 60.0 ± 7.2% at 10 μg/ml Dpo71 and 58.2 ± 7.0% at 40 μg/ml. Therefore, 10 μg/ml Dpo71 was chosen for the evaluation of synergistic effects with 1 μg/ml colistin (1/2 MIC) in inhibiting biofilm formation. Colistin alone brought down the biomass to 43.5% ± 4.9%, and further reduced to 28.9% ± 3.1% when used in combination with Dpo71 (**Fig. 4B**). The biofilm was visualized by the LIVE/DEAD staining, in which live cells were stained with green fluorescence and dead cells with damaged membrane were stained red (**Fig. 4C**). Our results confirmed the capability of both Dpo71-alone and co-treatment with colistin in preventing *A. baumannii* biofilm formation.

**FIG 4.**
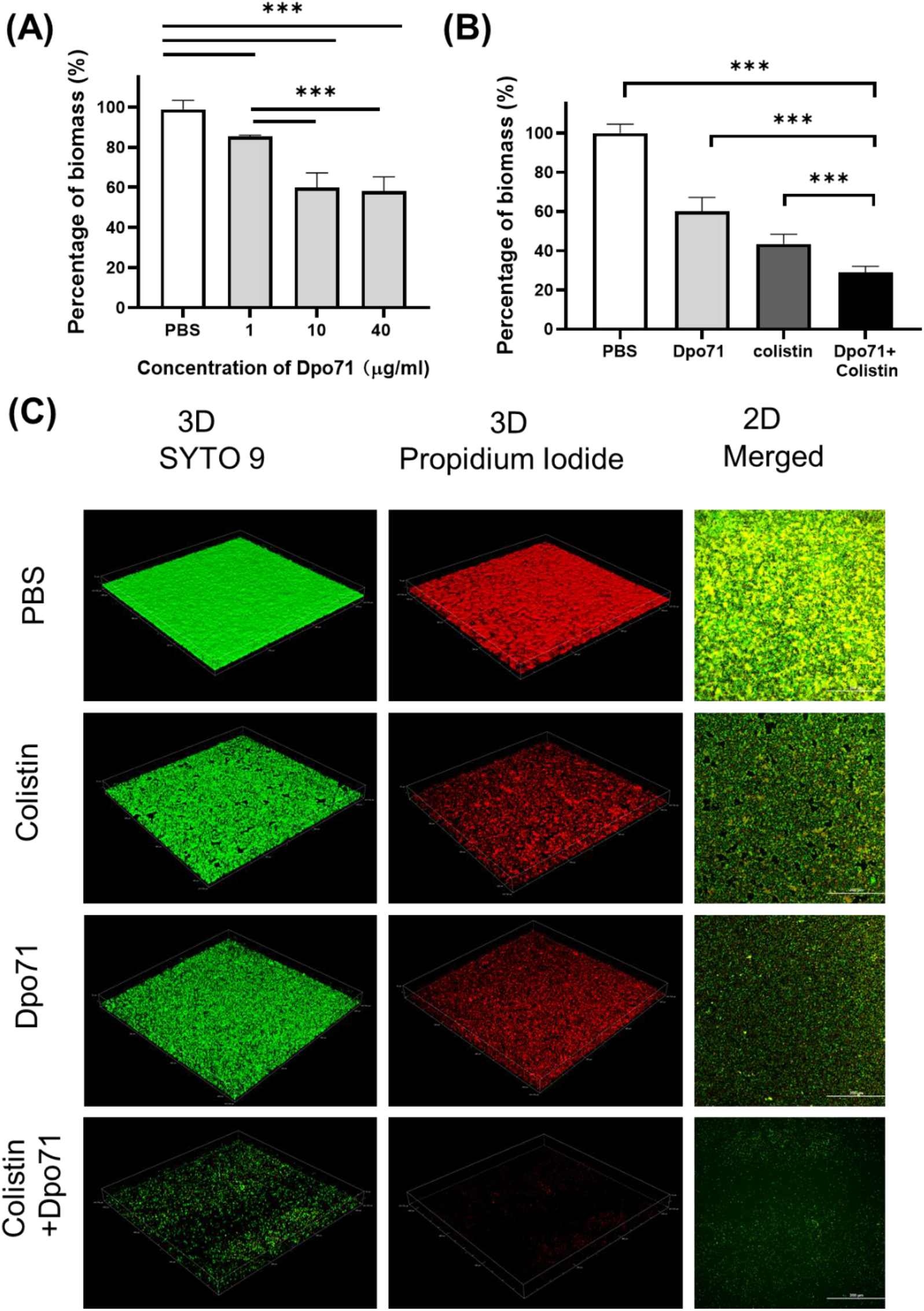
Dpo71 and colistin inhibited the biofilm formation. (A) Dpo71-alone inhibited the biofilm at a dose-dependent manner. (B) Dpo71 and colistin combination inhibited the biofilm formation. *A. baumannii* were incubated with PBS buffer; Dpo71; colistin (1 μg /ml); and their combination agents (Dpo71+colistin) in 96-well plates for 24 h, followed with crystal violet staining. Data are expressed as means ± SD (n =3) with **p<0.01, ***p<0.001 determined by Student’s t-test. (C) The Representative confocal fluorescence microscopic images of LIVE/DEAD stained *A. baumannii* biofilm (Scale bar, 200 μm).

We next assessed whether Dpo71 or its combination with an antibiotic can remove pre-formed biofilm. According to the CV staining assay, the biomass could be disrupted by a single treatment of Dpo71 (10 µg/ml) or colistin (4 µg/ml, 2× MIC) to around 60% residual biomass of the PBS control (**Fig. 5A**). The combined treatment could further reduce the residual biomass to 41.5% ± 6.6%. We reached the same conclusion when we compared the number of viable bacterial cells in the dispersed biofilms (**Fig. 5B**). In the absence of Dpo71, colistin was inefficient in killing bacteria embedded in the biofilm (< 0.5 log reduction), whereas with the help of Dpo71, more than 90% of the bacterial cells in biofilm were killed (from 7.3 ± 0.1 to 6.2 ± 0.2 in log scale). In the LIVE/DEAD viability assay, the pre-formed biofilm network was dismantled by Dpo71 (**Fig. 5C**). These results show that Dpo71 can efficiently disrupt per-formed biofilm.

**FIG 5.**
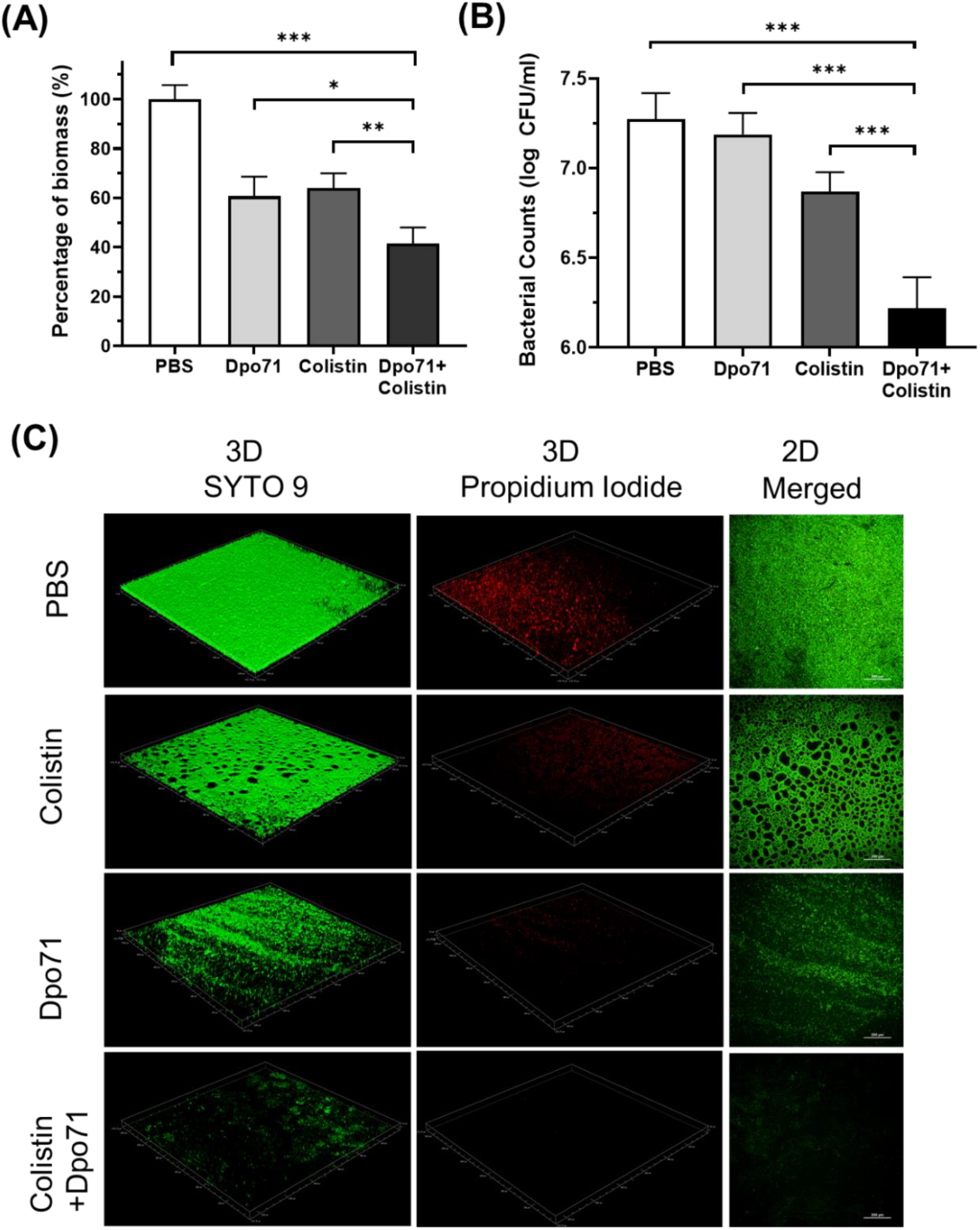
Dpo71 and colistin disrupted the pre-formed biofilm. The residual biofilm was assessed by (A) crystal violet staining and (B) *A. baumannii* bacterial counts. Briefly *A. baumannii s*train was grown on 96-well plates for 24 h for biofilm formation first, and then the biofilm was treated with PBS buffer, Dpo71 (10 μg /ml), colistin (4 μg /ml) or Dpo71+colistin for 24 h, followed with crystal violet staining and bacterial counting. Data are expressed as means ± SD (n = 3). *p<0.05, **p<0.01, ***p<0.001, Student’s t-test. (C) Representative confocal fluorescence microscopic images of live/dead stained *A. baumannii* biofilm (Scale bar, 200 μm). Confocal dish was used instead of the 96-well plate here.

### Antibacterial activity in a *Galleria mellonella* infection model

We next evaluated the *in vivo* efficacy of Dpo71 and colistin in combating bacterial infections in a *Galleria mellonella* infection model (**Fig. 6A**). In the control group, approximately 70% of the *G. mellonella* died within 18 h and the death rate increased to 90% at 48 h (**Fig. 6B**). The survival rate of the colistin-treated group increased to 50% after 24 h post-infection, and around 30% endured to the end of the monitoring period (72 h post-infection). Although depolymerase itself is not bactericidal, the Dpo71-alone treatment was found to be effective in rescuing the infected worms with 40% of the *G. mellonella* survived for 72 h. The combination treatment increased the survival rate of the infected worms to 80% till the end of the monitoring period, significantly higher than the monotherapy groups (**p < 0.01, log-rank test).

**FIG 6.**
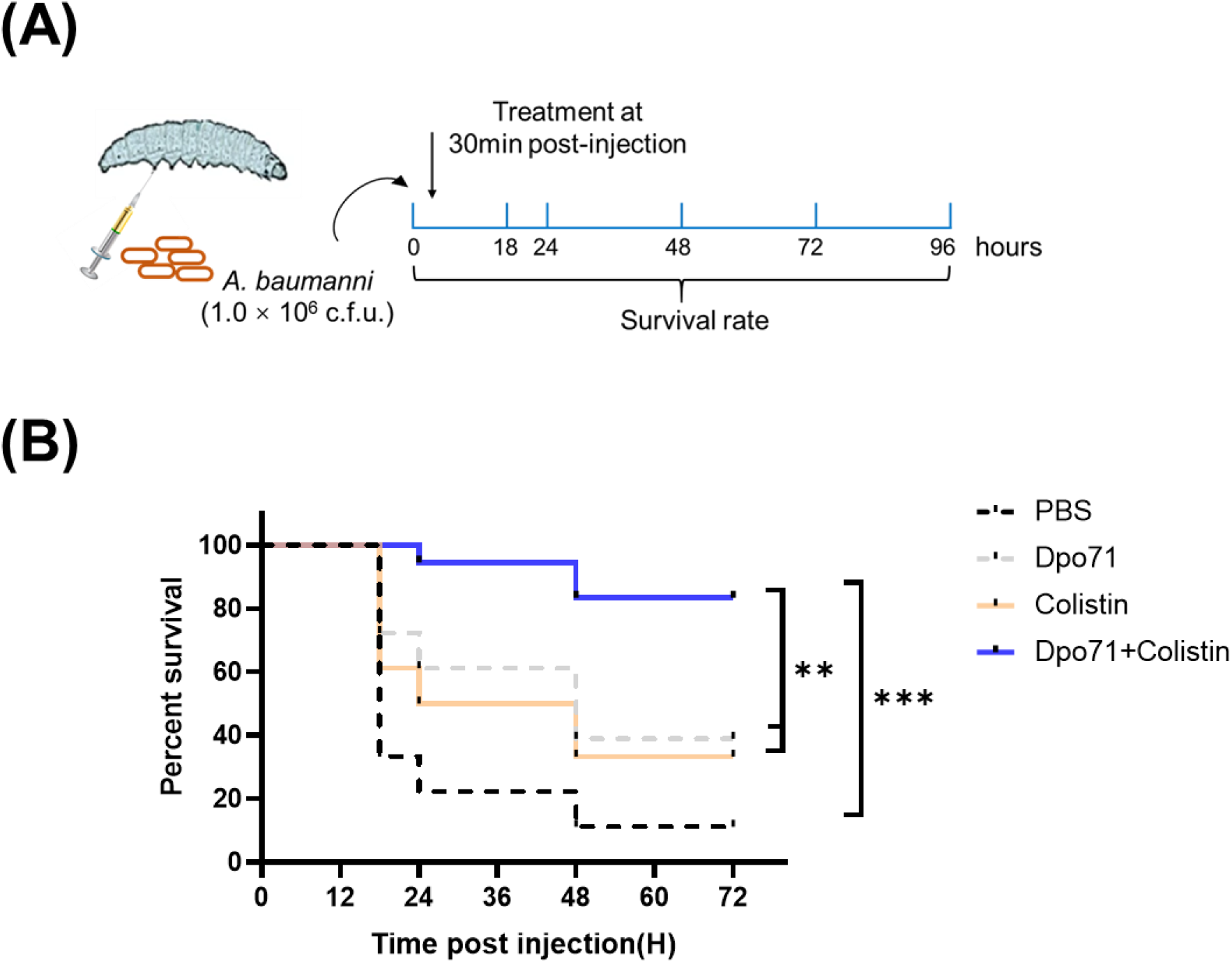
Antivirulent activity in the *Galleria mellonella* infection model. (A) Scheme of the experimental protocol for the *G. mellonella*. (B) Survival curves for *G. mellonella* infected with 10^6^ CFU *A. baumannii* then followed by the injection of PBS buffer (control group); 5 μg Dpo71; 1 μg colistin; or 1 μg colistin + 5μg Dpo71 (treatment group). (n=18, **p<0.01, *** p<0.001, Kaplan-Meier survival analysis with log-rank test).

## DISCUSSION

*A. baumannii*, one of most alarming nosocomial G-ve pathogens, has drawn significant attention in clinical settings due to its exceptional ability to acquire resistance to the commonly used antibiotics [40]. Although the knowledge on the mechanisms involved in the pathogenicity of *A. baumannii* is still limited, the production of bacterial capsular polysaccharides has been regarded as an important virulence factor, conferring its intrinsic resistance to peptide antibiotics and protecting it from host immune attack [41‒43]. *A. baumannii* has previously shown a much higher biofilm formation rate (80∼91%) compared with other species (5-24%) [44]. Its excellent capability of forming biofilms also contributes to the bacterial pathogenicity and resistance toward antibiotics. As CPS and EPS (a major component of the biofilms) are the substrates of phage encoded depolymerases, the application of recombinant depolymerases has received compelling interest as novel antivirulence agents to control multidrug-resistant infections [13,14]. A few depolymerases encoded by *A. baumannii* phage have been identified in recent years with demonstrated *in vivo* efficacy [28, 29, 32, 45, 46]. However, the synergistic effects of the combination of depolymerases and the SOC antibiotics in controlling infections caused by *A. baumannii* have never been attempted.

In this study, depolymerase Dpo71, derived from an *A. baumannii* phage, IME-AB2 was found to remain activity at a pH range of 4−8 and have a Tm of 58.5°C, suggesting it can be used under most physiological conditions. The robust nature of depolymerase was consistent with other reports [30, 35]. Importantly, it is the first report that demonstrates the excellent storage stability of a depolymerase with no noticeable activity loss for at least 6 months storing at 4 °C. This would offer great advantages in developing depolymerases as commercially viable antibacterial agents. The purified recombinant Dpo71 effectively decapsulates the host bacteria of its parent phage (AB#1 and AB#2) and re-sensitises them to serum killing in a serum ratio-dependent manner (**Fig. 3B**). The depolymerase treatment was largely limited for systemic infections because bacterial killing required the aid from the host immune attack such as complement-mediate killing. In the present study, the Dpo71 treated bacteria were significantly reduced in the presence of 5% serum ratio (4 log killing from a 10^8^ CFU/ml), representing the possibility of applying this depolymerase beyond systemic infection to environments with a low serum level, like lung infections. To the best of our knowledge, this was the first report showing that complete bacterial eradication could be achieved with depolymerase and serum (at 50% volume ratio) treatment. In previous studies [15, 21, 31, 35, 36, 46], a 50% serum (complement) ratio could only kill 2−5 log of the depolymerase treated bacteria and no further killing was noted when the serum (complement) ratio increased to 75%. Liu *et al*. postulated the incomplete bacteria killing was due to the emergence of the resistant phenotypes [31]. Therefore, the refractory to resistance development towards the depolymerase treatment was assessed and compared with the parent phage treatment (**Table 1**). While the bacteria developed phage resistance after 24-h co-incubation, they remained sensitive to the Dpo71 depolymerase. The sensitivity of the bacteria incubating with Dpo71 and 5% serum was also examined. Dpo71 could still yield a clear halo spot on the treated bacteria lawn (Data not shown), suggesting they were still sensitive to the depolymerase. It is mainly because depolymerases do not directly kill the bacteria during the antibacterial treatment, reducing the impetus for bacteria to evolve mechanisms against the depolymerases.

Biofilm formation is one of the major contributors for the chronicity of *A. baumannii* infections and their increased antibiotic resistance [44]. As the EPS can account for 80−90% of the biofilm matrix, the ability of phage in eradicating biofilms was reported to be accounted by the action of their tailspike depolymerases degrading the EPS, facilitating their diffusion through the dispersed biofilms to get access to the underneath bacteria [47]. The effectiveness of recombinant depolymerases in preventing biofilm formation and disrupting the established biofilms has also been studied. The susceptibility of biofilms to phage depolymerase treatments varied, depending on the bacterial strains and the activity of depolymerases. In most reported antibiofilm studies, depolymerases were able to cause a 10% – 40 % biofilm reduction compared with the untreated controls in a dose-dependent manner [18‒20, 25, 29]. However, there were also reports showing depolymerases were ineffective in dispersing the biofilms, though they were capable of decapsulating bacterial CPS [26]. Overall, the depolymerase treatment were unable to completely inhibit or remove biofiolms and the number of viable bacterial counts in the biofilms were similar to the untreated controls [18‒20, 25], with a few exceptions [24, 29]. These suggested that using depolymerases as a stand-alone treatment might not be sufficient in controlling infections associated with biofilms. Impairing drug diffusion (subdiffusion) within the biofilm matrix is a major contributor to the sub-optimal treatment to biofilm related infections [48]. Improving the antibiotics penetration into the biofilm matrix may hold the key to better clinical outcomes. In the present study, Dpo71 demonstrated moderate biofilm inhibition and removal capacities, both around 40% reduction compared with the PBS control, at an optimal concentration of 10 µg/ml (**Figs. 4 and 5**). In consistent with most previous studies, Dpo71 could effectively disperse the biofilms but failed to reduce the viable bacterial counts in the biofilms and this was also visually reflected in the LIVE/DEAD confocal images. Combination treatment with colistin, which is the only antibiotics that the MDR-AB2 strain susceptible to, was studied. The residual biomass and the number of viable bacterial counts within the biofilms were both significantly reduced compared with the depolymerase-alone and colistin-alone treatments, confirming the positive effect of depolymerase-antibiotic combination treatment noted in other bacterial species [23‒25]. Previously, Dunsing *et al*. [47] have proved that treating established *Pantoea stewartii* biofilms with phage tailspike proteins could rapidly restore unhindered diffusion of nanoparticles. Therefore, the improved antibiofilm ability with the combined Dpo71 and colistin treatment noted here was likely attributed to the improved colistin penetration within the biofilm matrix after the EPS depolymerization by the Dpo71. Overall, our data support the depolymerase and antibiotic combination as a promising alternative treatment strategies in managing biofilm associated infections caused by *A. baumannii*.

The *G. mellonella* infection model was first developed to study the bacterial pathogenicity by Peleg *et al*. [49] and has emerged as a valuable inset model to evaluate the effectiveness of novel antibacterial reagents [50, 51]. Several reasons make it a popular model: the larvae (1) can survive at 37 °C to mimic the physiological condition of human; (2) have fast reproduction time to allow high-throughput of experiments compared with mammalian systems; (3) have a semi-complex cellular and humoral innate immunity, which shares remarkable similarities with mammals, but no adaptive immune response to interfere with the therapeutic outcome [52]. Importantly, the *G. mellonella* model does not require ethical approval to provide informed data in reducing the number of mammals used for further identification/confirmation of the potential lead compounds. Liu *et al*. first evaluated the *in vivo* efficacy of Dpo48 identified from an *A. baumannii* phage (IME200) using a *G. mellonella* infection model [45]. They showed that the Dpo48 treated *G. mellonella* had a higher survival rate (10‒30%) than the untreated group at all the time points throughout the study period (72 h). Although the Dpo48 treatment outcome was not particularly profound in the *G. mellonella* infection model, they showed that the Dpo48 could significantly reduce the bacterial load 6 h post-treatment and rescue 100% of the infected mice (both normal and immunocompromised) from fatal sepsis. They attributed the difference in the insect and mammal infection models to the simpler innate immune response of insects. Nonetheless, their results confirmed that the *G. mellonella* infection model is sufficient to predict the antivirulent capacity of depolymerase and their ability to control *A. baumannii* infections in mammals. As shown in **Fig. 6B**, the survival rate of infected *G. mellonella* treated by Dpo71 or colistin monotherapy were around 40%, but the survival rate of those treated by the combination of Dpo71 and colistin could be significantly enhanced to 80%. The results confirmed that Dpo71 was effective in reducing the virulence level of MDR-AB2 *in vivo* and prolonged the survival time of the infected worms as demonstrated in Liu *et al*. [45]. Moreover, the adjuvant effect of depolymerase to colistin was demonstrated *in vivo*, in consistent with the *in vitro* biofilm experiments. These results warrant further studies on assessing the potential of the combination in treating biofilm-associated infections in mammals.

While promising effects on the use of depolymerase as antivirulence agents have been demonstrated *in vivo*, knowledge on the exact mechanisms of *A. baumannii* CPS cleavage by phage depolymerases are still largely missing [53]. In addition, depolymerases also present high specificity toward a narrow range of target polysaccharides (specific capsular type of the bacteria). In some cases, depolymerases might only be active against a subset of bacteria of their parent phage which are already specific to a small set of bacteria strains [18, 29, 36]. Such narrow host spectrum would greatly limit the wider therapeutic application of depolymerases. Further work on elucidating mechanisms of action of depolymerases would allow protein engineering to extend their host range and activity, facilitating their application as stand-alone treatment or as adjuvant with SOC antibiotics.

## CONCLUSION

In summary, phage tailspike proteins with depolymerase activity are promising antivirulence agents to re-sensitize *A. baumannii*, even the drug-resistant strains, to host immune attack. The identified Dpo71 depolymerase was found to effectively degrade bacterial capsules with excellent stability at various pH and upon storage. The refractory to resistance development toward depolymerase treatment have also made it attractive alternative weapon to control *A. baumannii* infections. In addition, Dop71 alone can be utilized to prevent and remove *A. baumannii* biofilms. The combination of Dpo71 and colistin was further demonstrated significantly enhanced antibiofilm activity compared with the monotherapies. Furthermore, this depolymerase was able to enhance the colistin antibacterial activity *in vivo*, markedly improving the survival rate of infected *G. mellonella*. As carbapenem-resistant *A. baumannii* has been ranked as the number one priority pathogen by the WHO and there are no antibiotics which have reached the advanced stage in the development pipeline to target this superbug, depolymerases as a stand-alone treatment or adjuvant to antibiotics may represent promising treatment strategies in controlling multidrug-resistant *A. baumannii* infections.

## MATERIALS AND METHODS

### Bacterial strains and culture condition

All bacterial strains used in this study are listed in **Table S1**. A multidrug-resistant strain of *A. baumannii*, MDR-AB2, isolated from the sputum samples of a patient with pneumonia at PLA Hospital 307 was supplied by the Beijing Institute of Microbiology and Epidemiology [34]. All the bacterial strains were grown in Nutrient Broth (NB) medium at 37 °C.

### Plasmid construction

The plasmid was constructed using standard cloning methods. Genes encoded Dpo71 (Protein id: YP_009592222.1) was synthesized by BGI (Shenzhen, China) and cloned into the pET28a plasmid using BamHI and XhoI site and the protein sequence was listed in supporting information (**Table S3**)

### Recombinant proteins expression and purification

#### Protein expression

The constructed plasmid was transformed into *E. coli* BL21 (DE3) cells and colonies were grown overnight at 37 °C in LB media supplemented with 50 μg/mL kanamycin. The start culture was grown overnight, and then was used to inoculate LB media supplemented with antibiotics at 1:100 ratio. The cell culture was grown at 37 °C to reach OD_600_ ∼0.6 before 0.25 mM IPTG was added to induce protein expression. After grown at 16 °C overnight, cells were harvested for protein purification.

#### Purification of depolymerase

The enzyme was purified by nickel affinity chromatography using HisTrap™ HP column (GE Healthcare). Briefly, harvested cells were re-suspended in lysis buffer containing 10 mM imidazole, 50 mM phosphate/300 mM sodium chloride (pH 8.0). The cell suspension was lysed by sonication and centrifuged. The supernatant was collected, filtered, and loaded into the column. The bound protein was eluted by imidazole gradient from 10 mM to 500 mM. Pure protein fractions eluted with imidazole gradient were collected and exchanged with PBS (pH 7.4). After purification, all proteins were flash frozen under liquid nitrogen and stored at -80 °C. Protein concentration was estimated using a Nano Drop (Thermo fisher, USA) using the extinction coefficient and molecular mass of corresponding lysins. Finally, the purity of the protein was analyzed by 12% SDS-PAGE.

### Circular dichroism (CD) spectroscopy

Far-UV CD spectroscopy is commonly used to analyse the secondary structures of proteins with a Jasco J810 CD spectrometer [35]. The spectrum measurement was performed with the Dpo71 of 0.30 mg/mL in PBS buffer (pH 7.4) using a wavelength range from 190 to 260 nm. Thermal denaturation with 1°C/min increments was also employed to measure the secondary structure unfolding at 215 nm, from 20 to 90°C. The melting curves were fitted into a Boltzmann sigmoidal function.

### Spot test assay

The depolymerase activity of Dpo71 was qualitatively assayed by a modified single-spot assay. In brief, 100 µL of MDR-AB2 overnight bacterial culture was added to 5 mL of molten soft nutrient agar (0.7%) and incubated at 37 °C for 3 h to form a bacterial lawn in plates. The purified enzyme was serially diluted, then 5 µL of each dilution (from 0.001 µg to 10 µg) was dropped onto an MDR-AB2 bacterial lawn for incubation at 37 °C overnight. The plates were monitored for the formation of semi-clear spots as a confirmation of the depolymerase activity.

### Extraction of bacterial surface polysaccharides

The extraction and purification of bacterial EPS was performed via a modified hot water-phenol method as described previously [36, 37]. Briefly, *A. baumannii* were cultured overnight in LB with 0.25% glucose. 1 mL culture was centrifuged (10000 rpm, 5 min) and resuspended in 200 µL of double distilled water (ddH2O). An equal volume of water-saturated phenol (pH 6.6; Thermo Fisher Scientific) was added to the bacterial suspension. The mixture was vortexed and incubated at 65 °C for 20 min, centrifuged at 10000 rpm for 10 min. Then the supernatant was extracted with chloroform to remove bacterial debris. The obtained EPS was lyophilized and stored at −20 °C.

### Quantification of depolymerase activity and alcian blue staining

The enzymatic activity of Dpo71 on bacterial surface polysaccharides was determined as described in Majkowska-Skrobek *et al*. [37] with minor modifications. The EPS powder of *A. baumannii* was resuspended in ddH2O (1 mg/mL), and mixed with Dpo71 (30 µg/mL) or deactivated Dpo71 (by heating at 90 °C for 15 min) to a final reaction volume of 200 µL. EPS or enzyme alone served as the controls. After 2 h incubation at 37 °C, cetylpyridinium chloride (CPC, Sigma-Aldrich) was added to the mixture at the final concentration of 5 mg/mL, which was further incubated at room temperature for 5 min. Absorbance was measured at 600 nm using a microplate reader (Multiskan Sky, Thermo Fisher). The experiment was performed in triplicate and repeated at least in two independent experiments.

The CPS detected by Alcian blue staining was performed as previously described [36, 37]. The treated samples were separated by a 10% SDS-PAGE. The gel was then washed with the fix/wash solution (25% ethanol, 10% acetic acid in water) and stained by 0.1% Alcian blue (Sigma-Aldrich) dissolved in the fix/wash solution for 15 min in the dark. CPS was visualized blue after the gel was destained overnight in the fix/wash solution.

### Influence of pH and storage time on the depolymerase activity

The EPS powder was dissolved in 100 mM citric acid-Na_2_HPO_4_ buffer (pH 3.0–8.0) or 100 mM Glycine-NaOH buffer (pH 9.0–10.0) to a final concentration of 1 mg/mL. The EPS solutions of *A. baumannii* were mixed with Dpo71 (30 µg/mL) to a final reaction volume of 200 µl, respectively. After 2 h incubation at 37 °C, the turbidity of residual EPS in various pH buffers was determined as described above. The effect of pH on the enzymatic activity was determined by this method. The storage stability of Dpo71 at 4 °C was determined by measuring the EPS degradation activity after 1, 3 and 6 months storage. All assays were performed in triplicate and repeated at least in two independent experiments.

### Serum killing assay

The serum killing assay was performed as previously described [35]. Logarithmic phase bacteria were prepared by inoculating overnight culture at 1:100 ratio in NB medium and shaking 180 rpm for 3-4 hours at 37 °C. Then the cells of each culture (AB#1, AB#2, AB#3 and AB#4) were harvested via centrifugation, washed, and resuspended in PBS, then adjusted to OD_600_ = 0.6. Human serum (Sigma, Shanghai, China) was mixed with (i) only bacteria or (ii) bacteria and Dpo71 mixture. The Dpo71 was fixed at final concentration of 10 µg/ml and volume ratio of human serum was at 50%. Experiments with heat-inactivated serum (at 56°C for 30 min) were served as controls. The mixtures were then incubated at 37 °C for 4 h for viable bacterial counting. Time-killing assays were also performed for the two sensitive strain (AB#1, AB#2) with the volume ratio of human serum varied from 1% to 50% and at 100 µg/ml Dpo71. Samples were withdrawn at 1, 3, 5 and 24 h for bacterial counting. The All assays were performed in triplicate and repeated at least in two independent experiments.

### Biofilm inhibition assay

*A. baumannii* strains were grown in NB medium overnight at 37 °C with continuous shaking 180 rpm. The overnight bacterial culture was diluted with fresh NB medium to a final density of 10^6^ CFU/ml. Then the diluted bacteria cultures were treated with PBS (control), Dpo71 (1 µg/mL, 10 µg/mL or 40 µg/mL), colistin (1 µg /mL) or their combination (Dpo71+ colistin) with a final volume of 150 µL / well at 37 °C for 24 h with gentle shaking (100 rpm). At the end of the incubation time, all medium were removed and the wells were stained with 200 µL 0.1% (w/v) crystal violet for 1 h. After staining, the crystal violet solution were removed and the wells were washed with 200 µL PBS for three times. Then, 200 µL of 70% ethanol was added to dissolve the crystal violet and 100 µL solution was transferred to a new plate for quantification of the residual biofilm biomass using a microplate reader (CLARIOstar, BMI Labtech, Germany) at 570 nm. All experiments were performed in three biological replicates and repeated at least in two independent times.

### Biofilm removal assay

*A. baumannii* strains were grown in NB medium overnight at 37 °C with continuous shaking 100 rpm. The overnight bacterial culture was diluted with fresh NB medium to a final density of OD_600_ = 0.2. To initiate the biofilm growth, diluted culture was aliquoted into a 96-well plate at 100 µL/well (Costar, Corning Incorporated, U. S. A.) and incubated at 37 °C for 24 h at 100 rpm. Biofilm was washed twice with PBS and treated with PBS (control), Dpo71 (10 µg /ml), colistin (4 µg /ml) or their combination (Dpo71+colistin) with a final volume of 150 µL / well at 37 °C for 24 h with gentle shaking (100 rpm). At the end of the incubation time, the residual biomass was quantified using crystal violet as described in the inhibition assay above. The antibiofilm activity was also evaluated by the counting the viable bacteria in the biofilm [54]. Briefly, the biofilm was grown and treated as above, then the wells were washed three times with PBS. Then the biofilm-containing wells were mixed fully with a pipetting device, making the biofilm cells become planktonic cells. Each sample was serially diluted and plated to for bacterial counting. All experiments were performed in triplicate and repeated at least in two independent times.

### Confocal laser scanning microscopy on biofilm removal

Confocal dish (NEST brand, China) was used instead of the 96-well plate for the antibiofilm study described above. Before microimaging, the biofilms were stained by LIVE/DEAD™ BacLight™ Bacterial Viability Kit (Thermo Fisher) for 60 mins following the manufacturer’s instructions. Then the biofilms were washed three times with PBS before the confocal laser scanning microscopy (Nikon, C2+ confocal, Japan) study. The excitation maximum and emission maximum of SYTO 9 is at 483 nm and 503 nm, respectively. The excitation maximum and emission maximum of propidium iodide is at 493 nm and 636 nm, respectively.

### Hemolysis assay

The effect of Dpo71 on the hemolysis of human red blood cells was performed using previously described methods with minor modifications [30, 31]. The human blood sample from a healthy donor was centrifuged at 1000 rpm for 10 min to remove the serum. The red blood cell pellets were washed with PBS (pH = 7.4) at least three times and then diluted to a concentration of 5% (volume ratio) with PBS. The Dpo71 (10 μg/ml,100 μg/ml and 500 μg/ml, final concentration) was added to the red blood cells and incubated at 37°C for 1 h, followed by centrifugation at 1000 rpm for 10 min. Then 100 µL supernatant was transferred to a 96-well microplate and topped up with another 100 µL of PBS to get a final volume of 200 µL. The erythrocytes in PBS and 0.1% Triton X-100 were served as negative and positive controls, respectively. The hemoglobin in supernatant was determined by measuring absorbance at 540 nm using a microplate reader (Multiskan Sky, Thermo Fisher). All experiments were performed in three biological replicates and repeated at least in two independent times.

### Cytotoxicity of Dpo71 against BEAS-2B cell

BEAS-2B (Human Normal Lung Epithelial Cells) cells were cultured in DMEM (Gibco) containing 10% FBS (Gibco) under standard conditions in a humidified incubator with 5% CO_2_ at 37 °C. The cytotoxic effect of the Dpo71 on BESA-2B cells was measured by Cell Counting Kit-8. The BEAS-2B cells were seeded at density of 10^4^ cells/well in a 96-well plate containing 200 µL of culture medium and incubated at 37 °C for 24 h. Next, the cells were incubated with Dpo71 for 12 h followed by incubating with 10 µL of WST-8 solution (Beyotime, Shanghai, China) for another 2 h at 37 °C. Absorbance was measured at a wavelength of 450 nm using a microplate reader (Multiskan Sky, Thermo Fisher). The PBS group was served as a negative control. All experiments were performed in three biological replicates and repeated at least in two independent times.

### Bacterial resistance assay

The development of phage-resistant and depolymerase-resistant bacterial variants was conducted as previously described [30] with minor modifications. Briefly, IME-AB2 phage (MOI of 10) or Dpo71 (100 μg/ml, final concentration) was incubated with ∼10^6^ CFU/ml of *A. baumannii* (MDR-AB2) in NB medium for 16 h (37 °C, 120 rpm). The cultures were plated to obtain isolated bacterial colonies. These were subcultured three times in agar plates to guarantee that the colonies were free of phage and Dpo71. Then 10 random colonies were picked to test the sensitivity toward both phage and Dpo71 using the spot test. Bacteria incubated with PBS was served as a control. Challenged bacteria were considered resistant to phage or Dpo71 when no plaque/inhibition halo was observed.

### *Galleria mellonella* infection model

The *G. mellonella* model was conducted following the procedures described by Peleg *et al*. with some minor modifications [49], and referring in other *G. mellonella* studies [50,51]. The *G. mellonella* larvae were acquired from WAGA company in Hong Kong and the injection was performed with a 10 μl SGE syringe (Sigma–Aldrich, Shanghai, China). To infect the *G. mellonella*, the larvae were first injected with 10^6^ CFU *A. baumannii* (MDR-AB2 strain) into the last left proleg. Then the PBS buffer (control group) or 5 μg Dpo71; 1 μg colistin; or 1 μg colistin + 5μg Dpo71 (treatment group), were injected into the last right proleg within 30 min. The *G. mellonella* were then incubated at 37°C and observed at 24 h intervals over 4 days. The *G. mellonella* which did not respond to physical stimuli were considered dead. Each group included 9 *G. mellonella* with individual experiments repeated two times (n = 18).

### Statistics

All experimental data are presented as means ± standard deviation (SD), and significance was determined using independent Student’s t tests and the one-way analysis of variance (ANOVA), assuming equal variance at a significance level of 0.05. Comparison of the survival rates of *G. mellonella* between groups was determined by Kaplan-Meier survival analysis with a log-rank test. All statistical analysis was performed using GraphPad Prism software.

### Data Availability Statement

All data for the paper are contained in the main text or SI Appendix.

## ACKNOWLEDGMENTS

The authors are thankful to Prof. Changqing Bai from the Fifth Medical Centre of Chinese PLA General Hospital, Beijing, China and Dr. Lin Zhang from Prince of Wales Hospital, Hong Kong for their kind donation of bacterial strains tested in the present study. This work was partially funded by the University Grants Committee of Hong Kong (ref. N_CUHK422/18 and ref. 24300619).

